# Identification of novel exosomal miRNAs and their role in diagnosis and prognosis of Triple Negative Breast Cancer

**DOI:** 10.1101/2025.06.13.659272

**Authors:** Ananya Choudhary, Satish S. Poojary, Priyanka Jain, Harit Chaturvedi, Bhudev C. Das

## Abstract

Triple-negative breast cancer (TNBC) is a clinically aggressive subtype with poor prognosis and limited treatment options. Exosomal microRNAs (miRNAs), encapsulated within secretory vesicles, have emerged as promising biomarkers for cancer detection and monitoring. In this study, we identify five novel exosomal miRNAs—hsa-miR-6803, hsa-miR-1180, hsa-miR-4728, hsa-miR-1915, and hsa-miR-940—that are consistently overexpressed in TNBC cells, stem-like subpopulations, and patient tumor tissues. Integrated analysis of public datasets and in vitro validation revealed that elevated expression of these miRNAs correlates with poor overall survival. Functional assays demonstrated that miR-1180 and miR-4728 significantly promote TNBC cell migration and invasion. These miRNAs also target critical oncogenic pathways, including Wnt, Notch, and EGFR. Their enrichment in exosomes highlights their translational potential as liquid biopsy-based biomarkers and therapeutic targets. This work is the first to link this miRNA panel to both TNBC tumorigenesis and stem-like cell biology, offering new insights into disease progression and potential strategies for personalized care.

## Introduction

Triple-negative breast cancer (TNBC) constitutes a highly aggressive and therapeutically challenging subtype of breast cancer, accounting for approximately 15–20% of all cases. Lacking expression of estrogen receptor (ER), progesterone receptor (PR), and HER2, TNBC is not amenable to endocrine or HER2-targeted therapies. As a result, treatment relies heavily on chemotherapy, which, despite initial effectiveness, is often followed by early recurrence, rapid disease progression, and poor overall survival. The absence of molecularly defined targets has driven an urgent need to discover novel biomarkers that can enable early detection, stratify risk, and identify new avenues for intervention

Within the TNBC landscape, cancer stem-like cells (CSCs) have been implicated as key contributors to treatment resistance, disease persistence, and metastatic relapse. These cells exhibit self-renewal, phenotypic plasticity, and the ability to evade cytotoxic therapies. Their survival following treatment may underlie minimal residual disease and tumor recurrence, making CSCs a critical focus for next-generation diagnostics and therapeutic strategies.

Exosome-derived microRNAs (miRNAs) have emerged as promising candidates for non-invasive biomarker development. Encased in lipid bilayer vesicles, these small non-coding RNAs remain stable in circulation and can reflect the transcriptional landscape of the tumor from which they originate. Exosomal miRNAs have been shown to mediate cell–cell communication, influence metastatic behaviour, and participate in therapy resistance. While several exosomal miRNAs have been explored in breast cancer broadly, the landscape of secretory miRNAs specifically enriched in TNBC and its CSC population remains poorly characterized. Moreover, the mechanistic significance of these miRNAs in TNBC biology has not been systematically addressed.

In this study, we present an integrative approach combining meta-analysis, experimental validation, and clinical correlation to identify a novel panel of exosomal miRNAs enriched in TNBC and TNBC-derived stem-like cells. We hypothesized that these secretory miRNAs not only reflect key molecular features of TNBC aggressiveness but also contribute to its invasive and therapy-resistant phenotype. Our findings aim to define exosomal miRNAs as functional biomarkers with translational potential for early detection, prognostication, and future therapeutic targeting in TNBC.

## Materials and Methods

### Identification and analysis of miRNA profiling datasets specific to TNBC

A search was conducted on PubMed and Web of Science electronic databases to find all the relevant literature studies on miRNA expression in TNBC. Referencing a method described formerly by Chen et.al.^81^, The search algorithms applied included”((microRNA OR micro RNA OR micro ribonucleic acid OR miRNA) AND ((breast carcinoma OR breast carcinomas OR breast cancer OR breast cancers OR triple negative breast cancer OR Triple Negative Breast cancers OR breast tumors) OR (TNBC carcinoma OR TNBC carcinomas OR TNBC OR adenocarcinoma of breast OR TNBC cancers OR TNBC)) AND (Humans [Mesh] AND English[lang]))”. Additionally, a total of 120 miRNA datasets were searched within the Gene Expression Omnibus (GEO) database (https://www.ncbi.nlm.nih.gov/geo/), with each dataset’s title, abstract, and full text thoroughly reviewed. The selection criteria included: (a) Original experimental studies that compare miRNA expression among different groups (TNBC vs. normal, TNBC vs. non-TNBC, and breast cancer vs. normal) using human samples, and (b) studies that report both upregulated and downregulated miRNAs, including specific cutoff parameters such as fold change and p-values. These specific inclusion criteria enabled the identification of all qualifying miRNA expression datasets. Comparably, we excluded datasets according to the criteria: (1) Any duplicated publications; (2) in-vitro or pre-clinical studies; (3) reviews, reports, editorials and (4) studies not characterised by miRNA expression analysis. A total of 53 studies on breast cancer were mined from the public domain and literature survey was conducted. The expression value of miRNA was calculated from each study. After selecting the datasets, data matrices were downloaded, and differential analyses were conducted using tools available on GEO. Various miRNA microarray platforms were employed, and uniquely expressed miRNAs from each dataset were annotated using miRbase. The fold change (FC) in miRNA expression was normalized by expressing it as log2FC to standardize the obtained miRNA expression values. Five miRNA showing high differential expression in breast cancer were selected for downstream experiments.

### miRNA target prediction

Potential miRNA–target interactions were predicted using established software tools, including TargetScan, miRTarbase, miRDB, and miRmap. The selection criteria included a prediction score of less than 0 for TargetScan, a cumulative score exceeding 95 for miRmap, and a prediction score range of 75 to 100 for miRDB, while all targets from miRTarbase were included for consideration. These miRNA sequences were utilized as input in conjunction with reference cDNA sequences in the miRanda tool.

### Gene ontology and enrichment analysis

Gene Ontology (GO) analysis is commonly utilized to assess the enrichment of differentially expressed genes (DEGs) in relation to biological processes, cellular components, and molecular functions. The candidate miRNA target genes were subjected to Gene Ontology (GO) and pathway enrichment analyses to elucidate their roles in critical biological pathways. Functional enrichment of predicted target genes for the selected five miRNAs was carried out using KEGG, Biocarta, Panther, and Reactome databases.

### Exosome isolation and analysis

Cells were grown in exosome-depleted, serum free media. Exosomes were isolated from cell free conditioned media collected at 48 hrs. by use of Exosome isolation kit (ExoCan Life Sciences, India). The exosome-containing supernatant was filtered through 0.2 μm membrane filters to remove particles exceeding 200 nm in size. Following filtration, the supernatant underwent centrifugation at 20,000×g for 40 minutes at 4°C to isolate the exosomes. The resulting pellet was then resuspended in 1× PBS for further processing to identify miRNA.

### Exosomal characterization (Physical properties)

EV characterization was conducted according to the International Society of Extracellular Vesicles guidelines. The physical properties of exosomes from cancer cell lines and stem cells were analysed using scanning electron microscopy (SEM) and transmission electron microscopy (TEM). A 100 µL aliquot of exosomes was placed on a formvar carbon-coated nickel grid for 1 hour. The grid was then cleaned over drops of 0.1 M sodium cacodylate (pH 7.6) and treated with a solution of 2% paraformaldehyde and 2.5% glutaraldehyde in the same buffer for 10 minutes. The grids were rinsed with M sodium cacodylate (pH 7.6), stained with 2% uranyl acetate for 15 minutes to enhance contrast, washed, treated with 0.4% uranyl acetate for 10 minutes, air-dried for 5 minutes, and then examined at 100 kV using a transmission electron microscope.

### Exosomal characterization (Western blotting)

The BCA Protein Assay Kit (Thermo Fisher Scientific) was employed to quantify exosomal proteins. Following electrophoresis on 4–15% gradient SDS-PAGE gels, 30 micrograms of protein were transferred to PVDF membranes, which were blocked using 5% bovine serum albumin (BSA) and subsequently incubated with specific primary antibodies, including CD63, CD81, and Calnexin (Biolegend, USA) at a dilution of 1:1000 for 24 hours at 4°C. Protein levels were assessed by probing with secondary antibodies against rabbit and mouse conjugated with horseradish peroxidase (HRP) at a dilution of 1:10,000, following three to five washing steps (10 minutes each). The bound complexes were detected using chemiluminescence methods (ECL; BioRad, USA), and images were acquired using the Amersham ChemiDoc Imaging System

### Exosomal miRNA isolation

Total RNA, including miRNA, was isolated from breast cancer cell lines in vitro utilizing the Trizol precipitation method. RNA quantity and quality were assessed using the NanoDrop 1000 spectrophotometer.

### Cell Culture

In this study, the human mammary epithelial cell line MCF-10A and TNBC cell lines MDA-MB-231 and MDA-MB-468, sourced from the American Type Culture Collection (ATCC, USA) were employed. Flow cytometric analysis revealed an enrichment of TNBC stem cells within the TNBC cell lines. The cells were cultured in Dulbecco’s Modified Eagle Medium (DMEM) supplemented with 10% exosome-depleted fetal bovine serum (FBS), 100 IU/mL penicillin, and 100 µg/mL streptomycin. Cultures were routinely maintained at 37°C in a 5% CO□ environment

### Immunocytochemical staining and immunofluorescence analysis

Cells were cultured on 0.13 mm thick coverslips (Corning® Cover glass, Corning Life Sciences) for 2 days, followed by washing with PBS and fixation in 1 mL of 100% methanol for 10 minutes. Permeabilization was then performed using 100% acetone for 30 seconds, followed by blocking with 1% BSA for 2 hours at RT. Primary antibodies (1-2 µg) were incubated with the samples overnight at a dilution of 1:1000, after which the samples were incubated with corresponding secondary antibodies. Finally, cells were mounted using VECTASHIELD mounting medium containing 0.5 µg/mL DAPI (VECTASHIELD; VectorLabs, CA)

### Western Blotting

For all western blot analyses of cell lines, proteins were extracted from cells that had reached approximately 80% confluence. The protein concentration in the cell lysates was determined using the BCA Protein Assay Kit (Thermo Fisher Scientific), and 30 µg of protein was loaded into each lane. Protein samples were subjected to SDS–PAGE (12%) and transferred onto PVDF membranes. The immunodetection protocol included transferring the proteins (15 V for 15 minutes per membrane) and blocking the membranes with a western blot blocking solution overnight. Following two washes with 1X TBST, membranes were incubated overnight at 4°C with primary antibodies against Sox2 and OCT4 (1:5000) and with GAPDH (1:5000) for 1 hour at room temperature. After five additional TBST washes, HRP-conjugated secondary antibodies (1:10000) were applied for 1 hour at room temperature, and detection was performed using ECL chemiluminescence.

### Culture of Tumorspheres

We assessed the capability of cell lines to generate spheres in an anchorage-independent suspension culture. Human breast adenocarcinoma cell lines MDA-MB-231 and MDA-MB-468 cell lines were cultured in DMEM (Gibco, ThermoFisher) containing 10% FBS (Gibco, ThermoFisher) and incubated at 37°C with 5% CO2 for 48 h. Following collection and washing, cells were resuspended in serum-free DMEM/F12 (Gibco, ThermoFisher) supplemented with 10 ng/ml fibroblast growth factor (FGF), 1% B27 and 20 ng/ml epidermal growth factor (EGF). At a density of up to 5000 cells/m, the cells were seeded in ultra-low attachment 6-well plates (Corning, USA) and incubated in a humidified environment with 5% CO2, set at 37°C for 4-6 days. Afterwards, the plates were examined for the growth of tumorspheres and measured with an inverted microscope. The tumorspheres were collected through mild centrifugation, followed by dissociation using Accutase (Sigma, USA) to produce individual cells, which were subsequently suspended in a serum-free medium to reform tumorspheres.. These tumorspheres were passaged every 5 days. Once the primary tumorspheres grew to a diameter of approximately 100µm, the samples were collected for downstream analysis.

### Flow cytometric analysis and CSC characterization

The expression of the molecular markers CD133□, CD44□, and CD24□ was evaluated by flow cytometry. Trypsin was used to carefully disperse tumorspheres into a single-cell solution after they were collected. Cells were labelled with anti-CD44-FITC, anti-CD133-PE, and anti-CD24 AlexaFluor antibodies (BD Biosciences, USA), incubated for 30 minutes in the dark at 4°C.The cells were analysed using a flow cytometer (BD FACS ARIA III). The acquisition was set for 10,000 events per sample. Data analysis was performed using the BD FACSDiva™ software (FACSDiva™, BD, USA). Cells were sorted based on the surface antigen expression.

### Side Population analysis

To isolate and identify SP and non-SP fractions, TNBC cells were treated with trypsin, then resuspended in pre-warmed DMEM containing 1% FBS.Dye Cycle Violet reagent (DCV) at a concentration of 5 µg/mL was added both in the presence and absence of verapamil (Sigma) and incubated at 37°C for 90 minutes with periodic shaking. Following incubation, the cells were washed with PBS containing 1% FBS, cold centrifuged, and resuspended in ice-cold sheath fluid (BD).Cells were preincubated with the ABCG2 inhibitor fumitremorgin-C(FTC) at a concentration of 10 μg/ml at 37°C for 30 minutes before DCV addition. Propidium iodide was added to the cells at a concentration of 1 µg/mL to differentiate viable cells. The Hoechst 33342 dye was excited at 357 nm, and its fluorescence was subsequently analysed using FACS AriaIII (BD Biosciences, San Diego, CA). The gating for forward and side scatter was stringent, ensuring that debris and non-viable cells were excluded from the analysis. Software used for analysis was BD FACSDiva™ (FACSDiva™, BD, USA).

### Immunofluorescence analysis

For tumorsphere immunostaining, cells were plated on glass coverslips (0.11 mm, Corning, USA) in DMEM with 10% FBS for 4 hours. Cells were then fixed with 4% paraformaldehyde and incubated with primary antibodies against SOX2 (mouse monoclonal IgG, Santa Cruz; 1:1000), OCT4 (mouse monoclonal IgG, BioLegend; 1:500), and ALDH1 (mouse monoclonal IgG, BioLegend; 1:500). Corresponding goat anti-mouse secondary antibodies conjugated with FITC, PE, and Cy3 were applied. Tumorspheres were incubated at 37°C for 60 minutes, followed by DAPI staining (Sigma) to visualize nuclei. Images were captured with a Zeiss fluorescence microscope, and processed using ZEN Blue and ZEN Black microscopy software (Zeiss)

### Transfection

TNBC cells were transfected with 50 nmol/L miRNA mimics of hsa-miR-1180 and hsa-miR-4728 using Lipofectamine 3000 reagent (Invitrogen). The cells were treated with the miRNA mimics and corresponding scramble controls (IDT Technologies) in Opti-MEM medium for 4 hours, then switched to standard growth medium as per the manufacturer’s instructions. Analysis was conducted 48 hours after transfection.

### Migration and Invasion assays

Matrigel (Corning, USA) was coated on top of the transwell chamber and the (serum starved) transfected cells were added in the upper chamber, seeded at 1 × 10□ /well. Subsequently, 500 μl of cell growth medium was added to the bottom chamber. Growth media serves as a chemoattractant. For 24 to 36 hours, the cells were incubated at 37°C. After the cells migrated to the lower chamber, they were fixed with 70% ethanol and stained using crystal violet. The quantity of migrated (stained) cells was measured by tallying the number of stained cells, and the mean cell count per field was computed for each well. Three replicate wells were utilized for every experiment, with representative images captured from randomly chosen fields in each well.

### Wound healing assay

TNBC cells (5 × 10□ cells/well) were seeded into 6-well plates and grown as a monolayer for 24–48 hours. Once cells reach a 80% confluence, a horizontal scratch is created in each well using a 200uL pipette tip. Detached cells were removed by rinsing with 500 μL PBS, followed by the addition of 500 μL of fresh medium. The plates were incubated for 12, 18, and 24 hours, and images were captured at each interval with an inverted microscope to observe scratch closure progress.

### Real-time PCR for miRNA expression analysis

Total RNA was isolated using TRIzol reagent (Invitrogen), with concentration measured at A260 and purity assessed via the A260/A280 ratio. RNA quality was evaluated using 2% agarose gels A total of 1 µg of RNA was reverse-transcribed into cDNA using random primers (Thermo), resulting in a final volume of 20 µL of cDNA. Subsequent amplification was performed via PCR for the miRNAs miR-940, 155, 6803, and 4728, with primer sets listed in Supplementary Table S1. RT-qPCR was conducted using SYBR Green chemistry,normalizing to U6 expression levels, and the ΔΔCt method was used to calculate normalized fold expression for the target miRNAs.

### Statistical analysis

All experimental data were quantified from three independent sets of experiments and expressed as mean ± standard deviation (SD). Statistical analyses were conducted using GraphPad Prism software. Differences between groups were assessed using Student’s t-test, one-way ANOVA, and two-way ANOVA, with significance levels determined as *p < 0.05, **p < 0.01, ***p < 0.001, and ****p < 0.0001. Additional study workflows are illustrated and included in supplementary file S1(Fig 1, 2).

**Fig 1 (a-e) :**
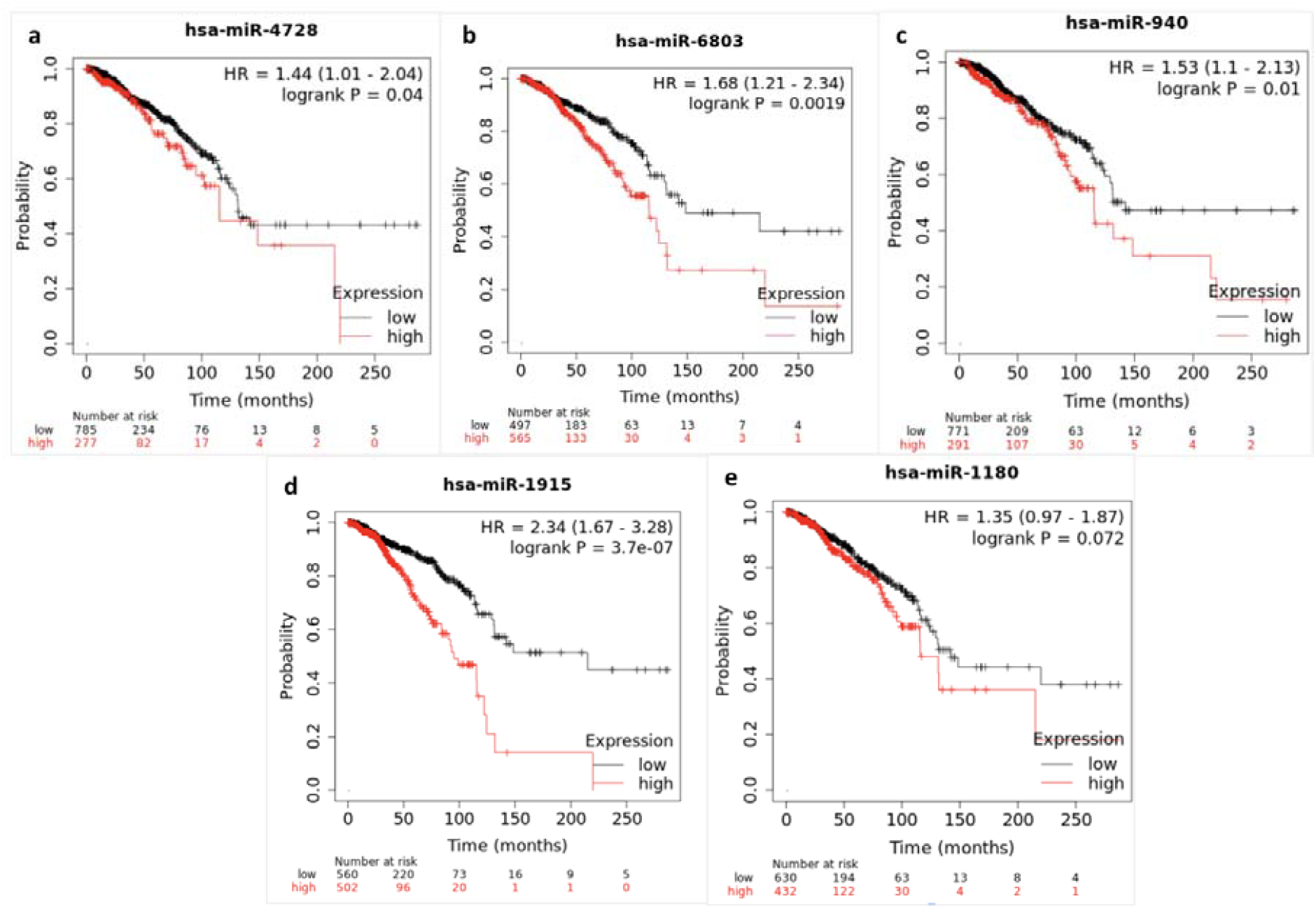
Expression of selected five miRNAs is correlated with overall survival (OS) in breast cancer using Kaplan Meier survival analysis (a-e) Graphs represent Kaplan-Meier curves for the 5 miRNAs and their impact on overall survival (OS). Increased miRNA expression is linked to poorer prognosis in breast cancer patients.

**Fig 2 (a-d):**
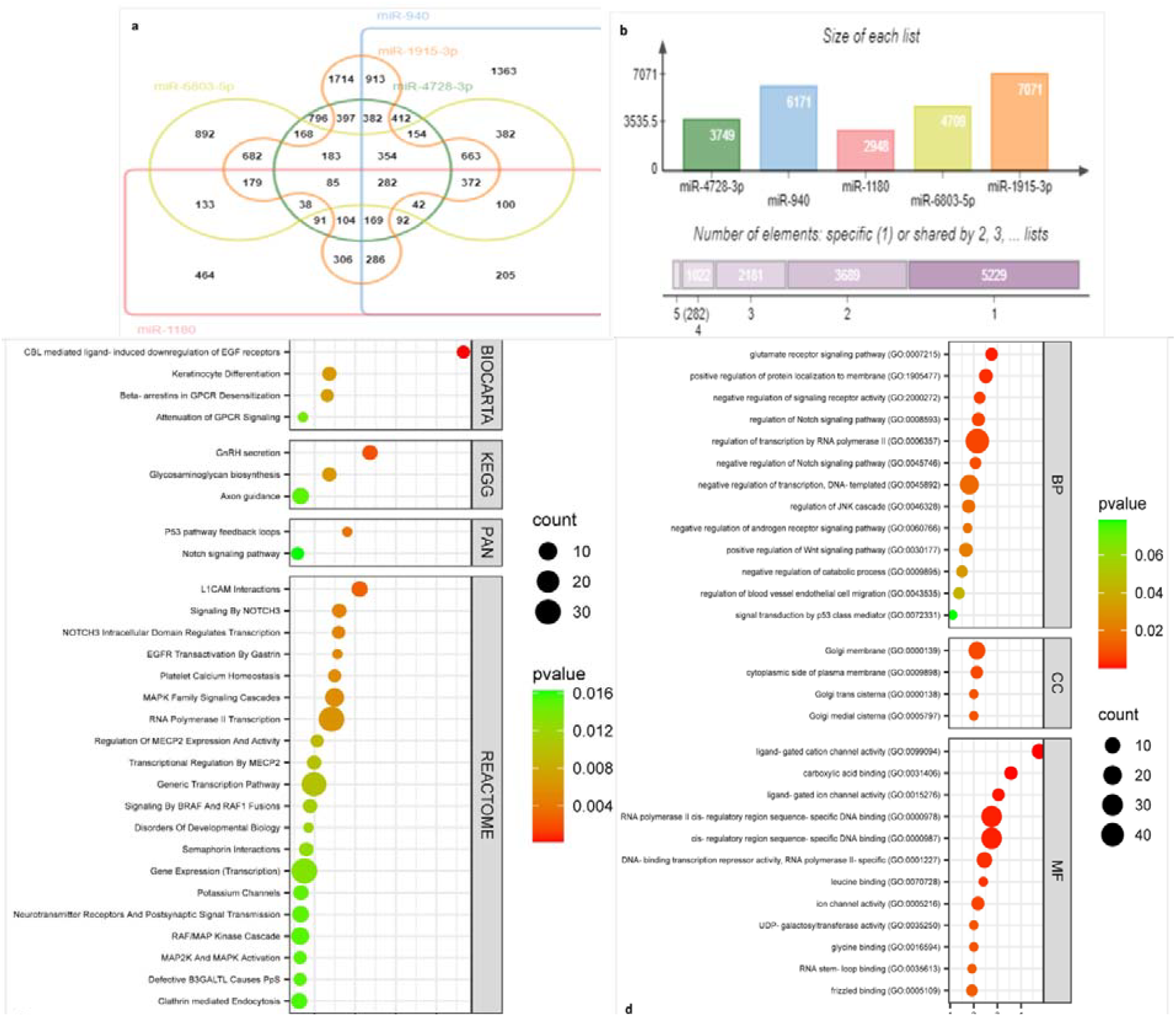
Intersection of predicted target genes from selected miRNA pool of miR-4728-3p, miR-940, miR-1180, miR-6803-5p and miR-1915-3p. (a-b): Indicates predicted target genes for differentially expressed miRNAs 4728-3p, 940, 1180 6803-3p, and 1915-3p. Functional enrichment analysis for predicted target genes of the selected five miRNA. (c-d) Pathway enrichment analysis was performed using the KEGG, Biocarta, Panther, and Reactome databases, in addition to Gene Ontology enrichment analysis.

## Results

### Identification of differentially expressed miRNAs by meta-analysis in breast cancer

For the meta-analysis, a total of 120 miRNA datasets related to breast cancer were obtained from the Gene Expression Omnibus (GEO) databases. Of these, a sum of 53 datasets on breast cancer were shortlisted and mined from the public domain based on various exclusion and inclusion criteria. Following the dataset selection, we downloaded data matrices and differential analyses were conducted using the GEO2R tool provided on the GEO platform. Across these datasets that reported multivariate analyses, expression values of miRNAs were calculated from each study. A set of five miRNAs showing high differential expression in invasive breast cancers was selected for downstream experiments. The predictive significance of the chosen miRNAs in breast cancer patients was validated by performing Kaplan-Meier analysis which demonstrated that all five miRNAs : hsa-miR-6803, hsa-miR-1180, hsa-miR-4728, hsa-miR-1915 and hsa-miR-940 were significantly associated with poor overall survival (OS). Survival analysis indicates increased expression of these miRNAs is correlated with poor prognosis for patients with breast cancer as presented in Figure 1 (a-e). While these miRNAs have shown poor prognosis across breast cancers, there is limited data to support the same in case of TNBCs as these miRNAs have been previously unreported in TNBCs and TNBCSCs. Given the prognostic promise and limited availability of data, we explored the prognostic potential of these miRNAs in TNBC.

### Target prediction and Functional annotation for target genes

To further identify their target genes of these five selected miRNAs, target prediction was performed using four electronic databases-TargetScan, miRTarBase and miRDB and miRmap. We created a Venn diagram to illustrate the intersections between five miRNA targets, summarizing common targets across breast cancers in the initial cohort (Fig 2a). The number of predicted targets for miR-4728-3p, miR-940, miR-1180, miR-6803-5p and miR-1915-3p was 3749, 6171, 2948, 4709 and 7071, respectively. (Fig 2b) Gene set enrichment analysis is widely used to evaluate the enrichment of differentially expressed genes (DEGs) in a range of biological processes, cellular components, and molecular functions. Analysis of DEGs using Gene Ontology revealed significant enrichment in processes such as endothelial cell migration, signalling pathways involving Wnt, NOTCH, EGFR, the JNK cascade, along with cell division and DNA replication. (Fig 2c-d). These results indicate the potential involvement of key signaling pathways and biological processes with our selected miRNAs, leading to tumor progression.

### Isolation and characterization of exosomes derived from TNBC cells

Exosomes were collected from the conditioned media of TNBC cell lines MDA-MD 231 and MDA-MB 468 after 48h. We performed scanning electron microscopy (SEM) and transmission electron microscopy (TEM) to examine cell culture-derived EVs. Further, Confocal microscopy was employed to qualitatively analyse localization of exosomes within TNBC cells. Exosomes were successfully identified as double-membrane vesicles through TEM Fig 3 (d,h)) and their size range was estimated to lie between 50-200 nm as indicated by SEM (Fig 3(c,g)). The exosome marker CD81 was successfully found localized in TNBC cells as indicated by Immunofluorescence (Fig 4 (a-c) and western blot Fig 4(d)

**Fig 3(a-h).**
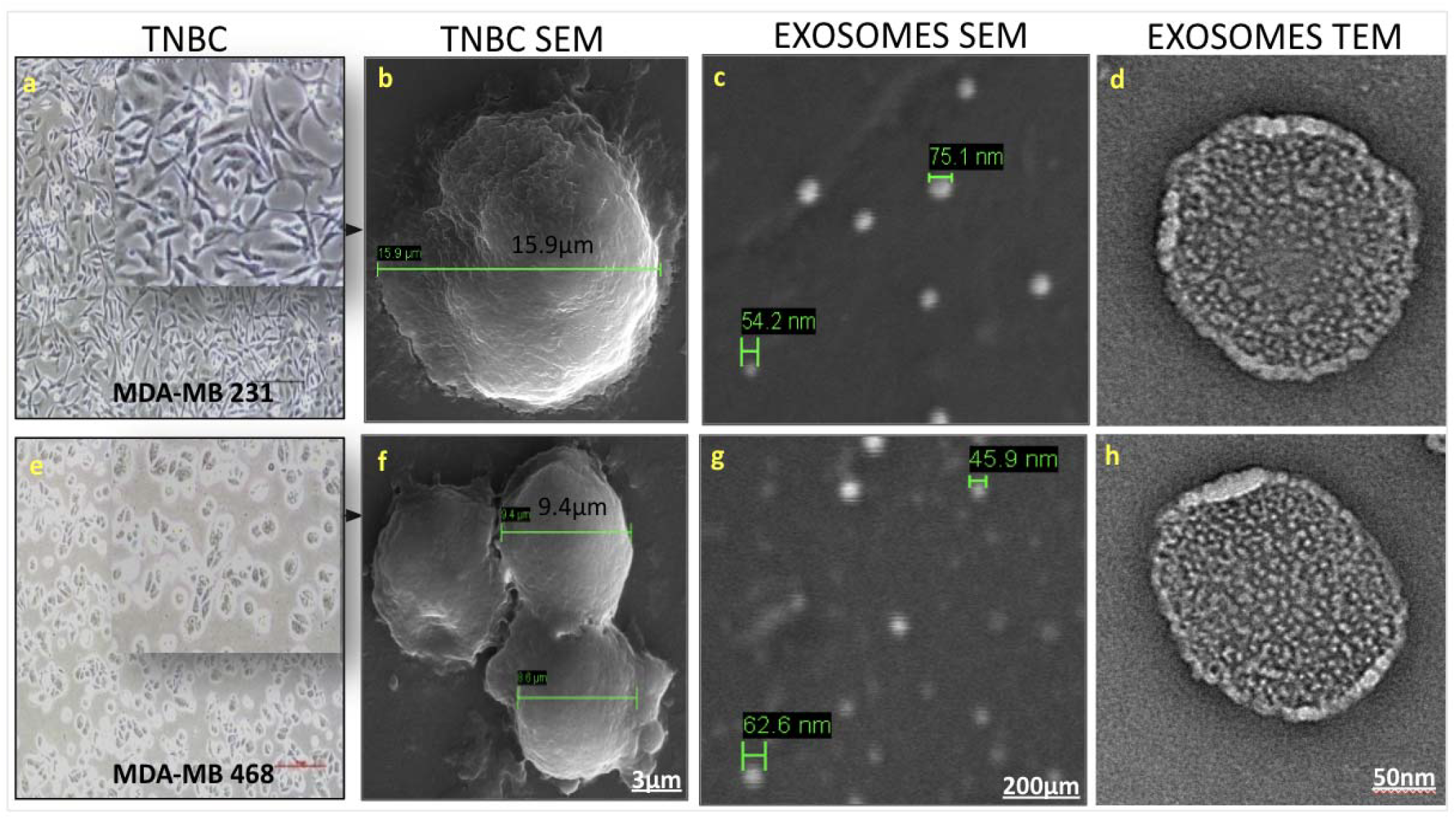
Characterization of TNBC exosomes using advanced microscopy techniques. Phase contrast images of TNBC cells MDA-MB 231 and MDA-MB-468 (a,e), Scanning electron microscopy images of TNBC cells (b,f) scale 3um and of TNBC derived exosomes (c,g) scale 200um. High resolution transmission electron microscopy (HRTEM) imaging of TNBC derived exosomes (d, h), scale 50 nm

**Fig 4 (a-c):**
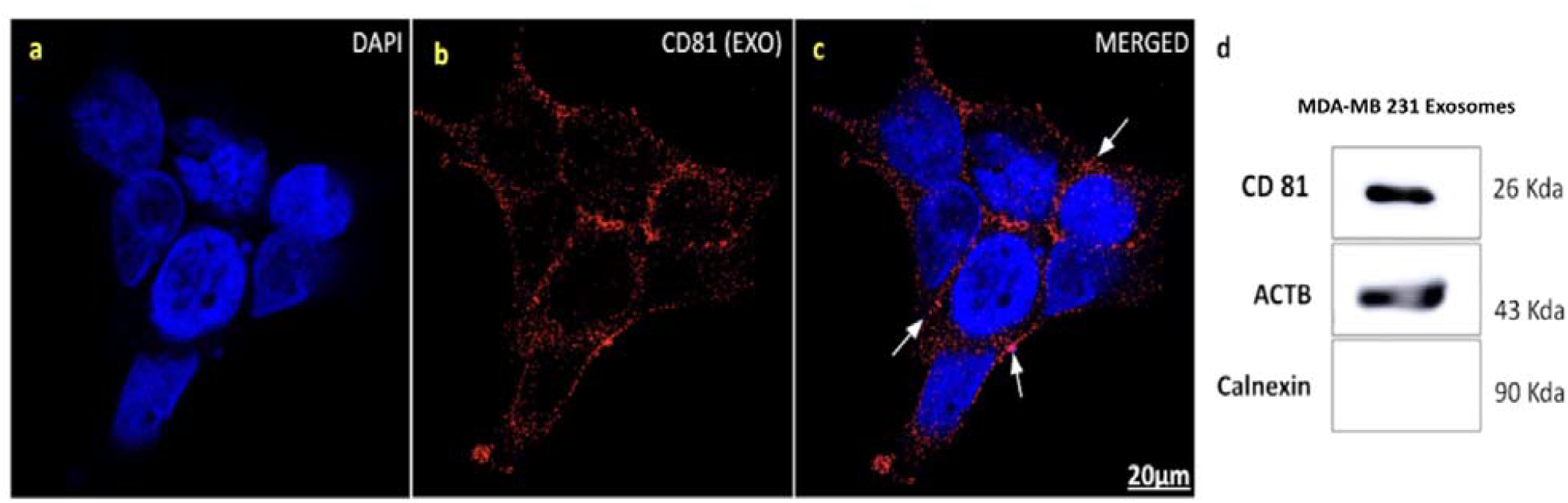
Representative Immunofluorescence images showing CD81 staining in TNBC cells (arrows indicate presence of exosomes). Scale bar 20um. Fig 4(d): Immunoblot showing expression of CD81 exosomal marker in TNBC cell line MDA-MB 231 derived exosome samples. Calnexin is used as a negative control while β-Actin is used as a positive control.

### Presence of cancer stem-like cells in TNBC

Using flow cytometric techniques, we identified the presence of stem-like cells in the TNBC population. These cells have the potential to generate 3D spheroids in culture, exhibiting stem-like properties. Our results exhibit increased expression of stemness markers SOX2, ALDH1 and ABCG2 in tumor-derived spheroids. This was confirmed through RTqPCR analysis (Fig 6G) and Immunoblotting (Fig 6H). Furthermore, expression of stemness marker genes in TNBCSCs was found to be two-fold higher compared to TNBC cells (Fig 6H).

Flow cytometric analysis was carried out in TNBC cell lines MDA-MB 231 and MDA-MB 468 to evaluate for expression of stemness marker CD133 and CD44 and identification of cancer stem cells. FACS analysis revealed a presence of a small population of SP cells (0.9-1.1%) that exhibit dye exclusion properties of cancer stem-like cells (Fig 5 (e-h)) The SP cells, which disappear in the presence of verapamil (f,h), are outlined and shown as a percentage of the total Cancer stem-like cell population. SP cells were subsequently grown in cell culture and developed tumorspheres as seen in representative phase contrast images (b,d). Further, marker-based selection of TNBC stem cells by FACS also revealed presence of small population (1-2%) of stem-like cells expressing markers CD133_D_, CD44_D_, and CD24_D_ (Fig 5 (i-l)). TNBC stem-like cells were characterized by immunostaining of tumorspheres with SOX2, ALDH1 and ABCG2 markers (as seen in Fig 6 (A-F) and immunoblotting to determine the expression level of ALDH1 (Fig 6G).

**Fig 5(a-l):**
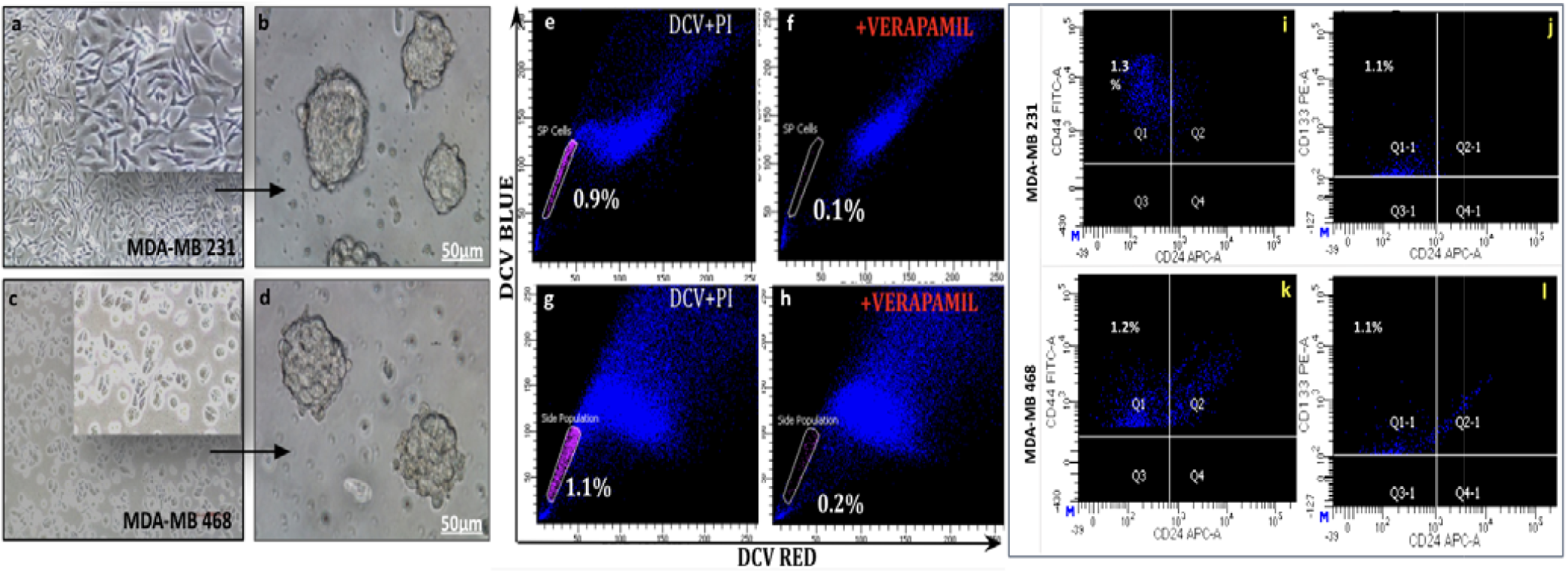
Isolation and characterisation of TNBC stem-like cells by FACS. Phase contrast imaging of TNBC cells and TNBC Stem-like cells (a-d). CSCs are observed as spheroids in culture (b,d). Flow cytometric analysis of TNBC cells using DCV based side-population assay (e-h). FACS analysis using marker-based selection methods expressing markers CD133□, CD44□, and CD24□ (i-l)

**Fig 6 (A-H).**
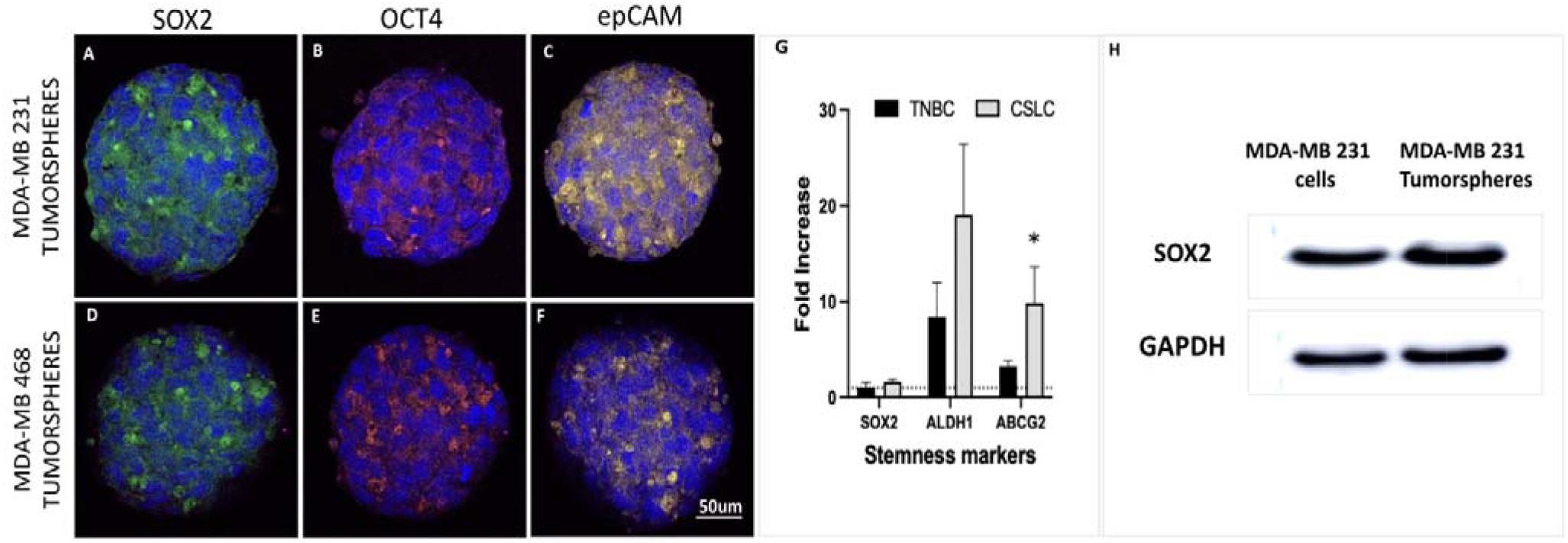
Characterization of TNBC derived stem-like cells. Immunofluorescence imaging of TNBC stem-like cells (A-F) Tumorspheres indicate presence of stemness markers SOX2, OCT4 and EpCAM. DAPI is used as a nuclear stain. SOX2 is tagged with FITC dye and thus emits green fluorescence, OCT4 is indicated in red (AlexaFluor 647) and EpCAM is marked by yellow dye (PE). Scale bar is 10um. Expression levels of stemness marker genes SOX2, ALDH1 and ABCG2 in TNBC and Cancer stem-like cells (CLSC) by qRT-PCR (G) Representative immunoblot showing expression levels of SOX2 in TNBC tumorspheres and cancer cells. GAPDH was used as positive control (H)

### Targeting hsa-miR 1180 and hsa-miR 4728 inhibits invasion and migration in vitro

To explore the role of miR1180 and miR-4728 on cell growth and proliferation in TNBC, we performed loss of function experiments by transfecting TNBC cells with anti-miR-1180 and anti-miR-4728 (24 h) in TNBC cells. As shown in Figure 7(A) in the wound healing assay, knockdown of miR-1180 and miR-4728 inhibited cell migration and proliferation when compared with WT conditions. Figure7(B) shows a drastic reduction in TNBC cell invasion abilities when miRNAs-1180, 4728 were silenced compared with the WT control. These results indicated the role of miR-1180 and miR-4728 in the cell proliferation, migration and invasion in TNBC cells.

**Fig7(A).**
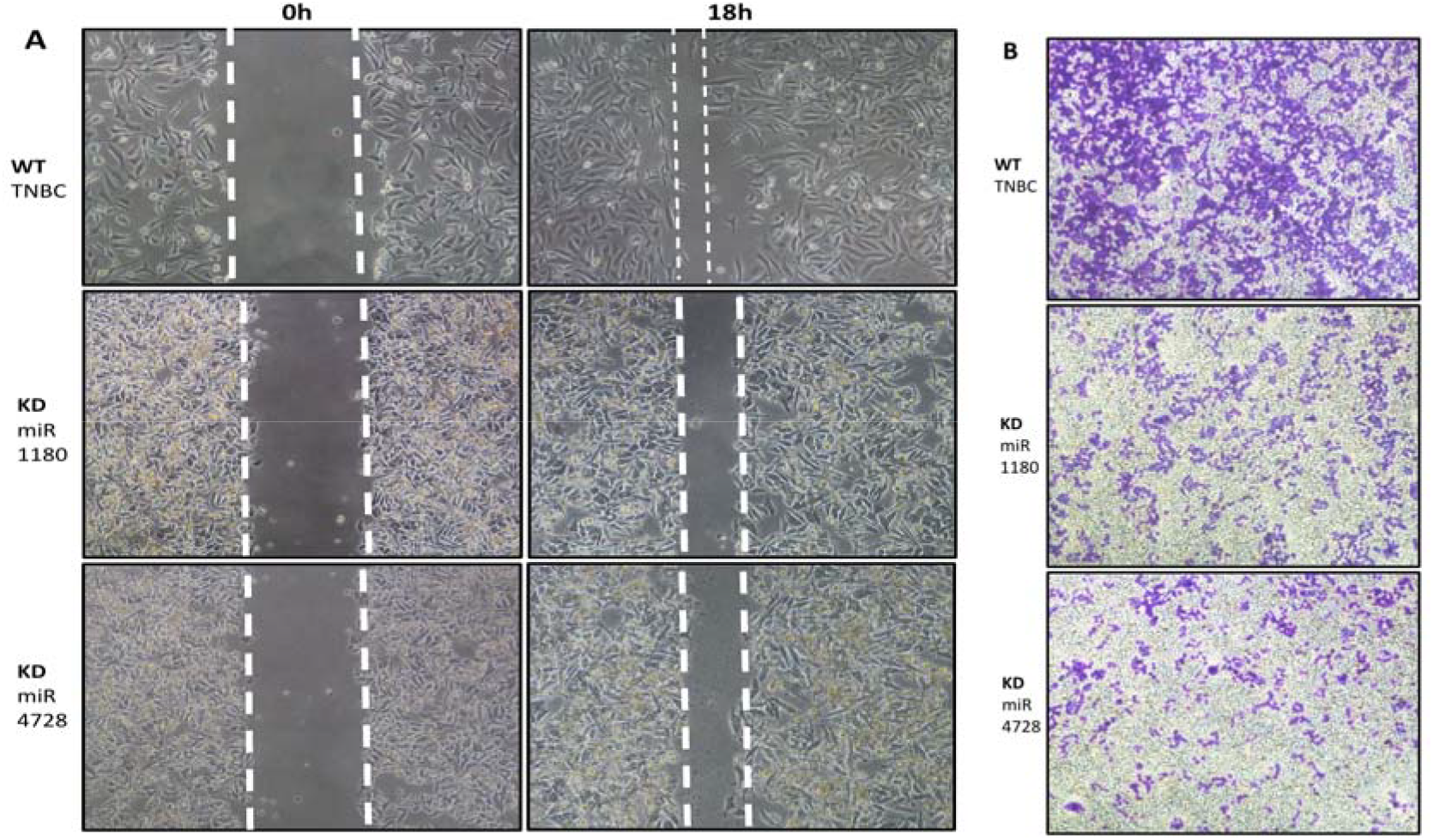
Wound healing assay of TNBC cells MDA-MB 231 (WT) and KD miR-1180 and miR-4728. Photographs were taken at intervals of 0h and 18h respectively. (B) Cell invasion assay using WT TNBC and KD of miR-1180, miR-4728. After 48h, cells were fixed with methanol and stained with crystal violet. Subsequently photographs were taken in phase contrast microscope. Transiently transfected cells with inhibition of miR-1180, miR-4728 show significant reduction in cell invasion in TNBC. These results reveal that miR-1180 and miR-4728 play a significant role in TNBC proliferation, migration and invasion.

### Exo-miRNAs consistently expressed in CSCs and breast cancer tissues

RT-qPCR was employed to assess the levels of oncogenic miRNAs in TNBC cells and stem cells in vitro (Fig 8 a, b, d, e). We found a significant upregulation of our target miRNAs miR 4728, miR 6803, miR-940, miR - 1915 and miR-1180 in TNBC cell lines MDA-MB 231, MDA-MB 468 in comparison to their corresponding CSCs. A similar expression pattern was observed in TNBC and TNBCSCs derived exosomes (Fig 8c and f). Our panel of five miRNAs-miR 6803, miR 1180, miR 4728, miR 1915 and miR 940 were found to be highly expressed in TNBC cells, TNBCSCs and enriched in circulating exosomes. Given their circulatory nature and consistent expression across TNBC cells and stem cells, these results indicate miRNAs miR 4728, miR 6803, miR-940, miR -1915 and miR-1180 may have a significant role to play in TNBC extracellular communications leading to disease progression or better prognosis if these miRs are targeted for therapy.

**Fig 8 (a-g):**
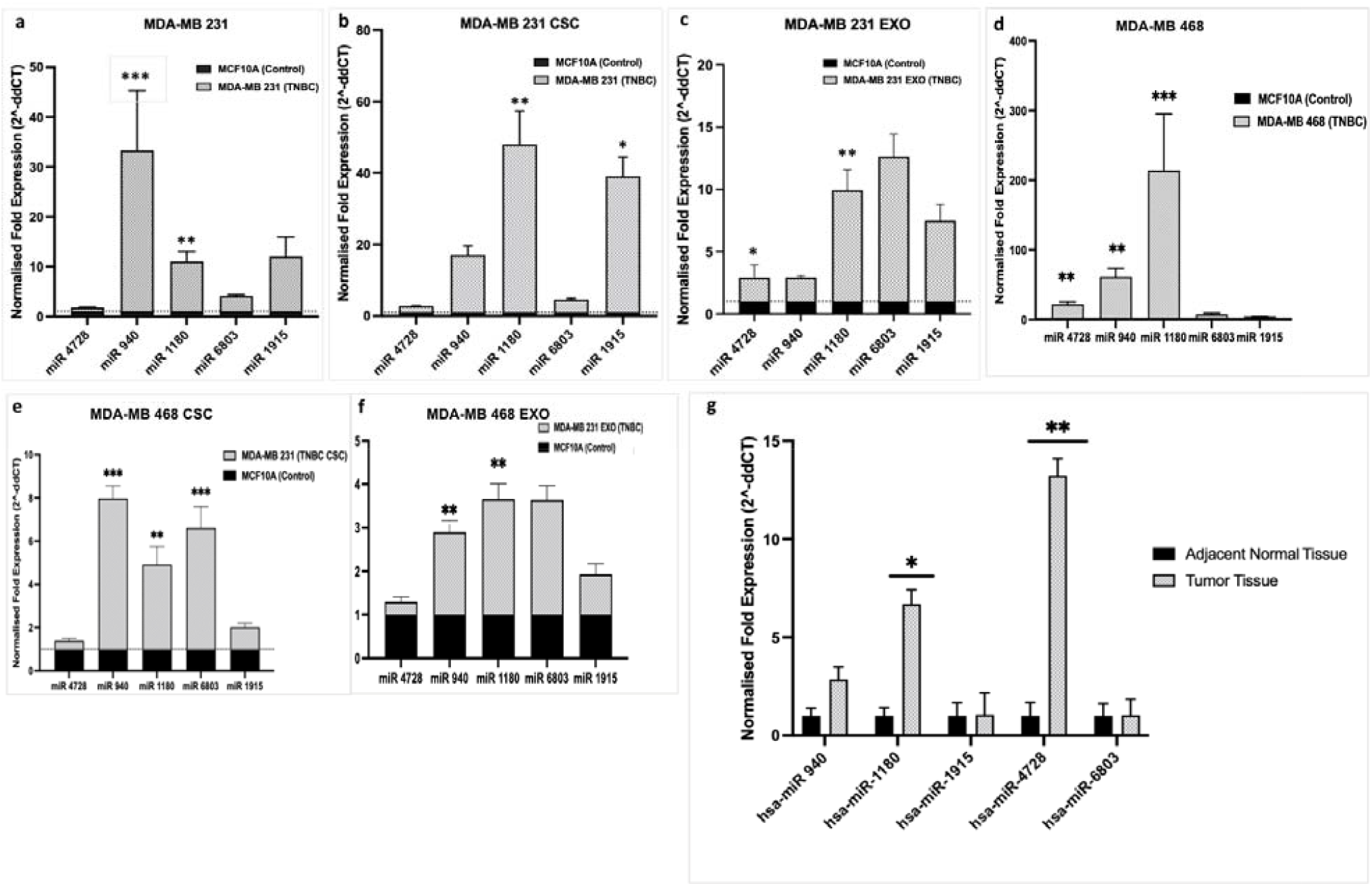
RT-PCR was employed to assess the levels of oncogenic miRNAs in TNBC cells, stem cells and tumor derived exosomes in-vitro. Expression analysis of miRNAs in TNBC cells (a,e). miRNAs were detected in TNBC exosomes (c, f). Selected miRNAs expressed in breast cancer tissues (n=15) as seen in 8(g). hsa-miR 1180 and hsa-miR 4728 are found to be highly upregulated in TNBC tumor tissue when compared with adjacent (normal) tissue

## Clinical Correlation

To clinically validate our findings, the presence of these miRNAs was evaluated in breast cancer tissues. Tissue biopsies were obtained from 15 TNBC patients, along with corresponding adjacent normal tissue samples for comparison. RT-qPCR analysis of resected tumor tissue RNA revealed consistent expression of these miRNAs in patients with TNBC (Fig 8g). Amongst the five miRNAs, hsa-miR 1180 and hsa-miR 4728 were found to be significantly upregulated in TNBC biopsies when compared with their adjacent (normal) tissues. Thus, our results demonstrated a positive correlation between the elevated expression of these miRNAs and the progression of TNBC. Second, the presence of hsa-miR 1180 and hsa-miR 4728 in circulatory tumor-derived vesicles and the detection of the same miRNAs in clinical tissue samples highlights their significance as potential biomarkers in TNBC.

## Discussion

### Summary of Findings

This study identifies a novel panel of five exosomal microRNAs—hsa-miR-6803, hsa-miR-1180, hsa-miR-4728, hsa-miR-1915, and hsa-miR-940—that are consistently overexpressed in triple-negative breast cancer (TNBC) cell lines, cancer stem-like cells (CSCs), and clinical tumor samples. Integrative meta-analysis, in vitro functional assays, and clinical correlation collectively demonstrate that these miRNAs are associated with poor prognosis and may serve as key mediators of TNBC aggressiveness. Notably, miR-1180 and miR-4728 were functionally validated to promote invasion and migration in TNBC cells, further supporting their role as drivers of tumor progression. Currently there are hardly any targeted therapies for combat this aggressive and debilitating disease. Through intercellular communication with the tumor microenvironment and potentiated by CSC population, TNBC cells often acquire treatment resistance leading to metastasis and chemoresistance^26,44–47^. Additionally, the molecular heterogeneity of TNBC as revealed by the presence of many molecular markers poses challenges to effective treatment.^48^ Several studies have outlined promising predictive and prognostic markers in breast cancer^49–51^. However, very few studies have been able to identify TNBC specific therapeutic biomarkers^52,53^. At present, there is no universal biomarker available for diagnosing and targeting TNBC, unlike other breast cancer subtypes, which have biomarkers such as HER2 and specific hormone receptors^49,50^ There is a critical need to identify reliable, non-invasive markers specific to TNBC for rapid screening, early diagnosis, risk assessment, and ultimately for effective treatment and management of breast cancer. This study highlights the clinical utility of specific exosomal miRNAs as non-invasive diagnostic and prognostic markers, as well as potential therapeutic targets.

### Biological Implications of Identified miRNA

The identified miRNAs appear to regulate pathways central to TNBC pathogenesis. Enrichment analyses revealed their involvement in oncogenic signalling cascades such as Wnt/β-catenin, Notch, and EGFR pathways. These signalling routes are known to maintain stemness, promote epithelial-to-mesenchymal transition (EMT), and enhance survival in CSCs. The persistent expression of these miRNAs in both TNBC cells and their exosomal cargo suggests a mechanism for reinforcing autocrine and paracrine signalling within the tumor microenvironment, thereby contributing to cellular plasticity and therapeutic resistance.

After comprehensive literature review and subsequent meta-analysis of 120 publicly available breast cancer datasets, 53 relevant datasets were identified based on various exclusion and inclusion criteria. Upon analysis of these datasets, a set of five, highly oncogenic miRNAs were found to be expressed in TNBCs : hsa-miR 6803, hsa-miR 1180, hsa-miR 4728, hsa-miR 1915 and hsa-miR 940. Though these miRNAs were often found in breast cancers, they have never been shown to be exclusively reported in TNBC, making them ideal candidates for therapeutic interventions. Additionally, since there was so far no evidence linking these oncomiRs with disease relapse, we were eager to explore their gene expression profile in TNBCSCs.

miRNA target prediction was carried out using four electronic databases :TargetScan, miRTarBase and miRDB and miRmap, which revealed 14 common targets between the two selected miRNAs - hsa-miR-1180 and hsa-miR-4728. PPI and RNA-protein association analysis of targets using the Search Tool for the Retrieval of Interacting Genes/Proteins (STRING) database (https://string-db.org) revealed amongst others, their role in epigenetic regulation of HATs, TP53 regulation of metabolic pathways and PPAR□ mediated gene expression in breast cancer (Fig 9)

**Fig 9:**
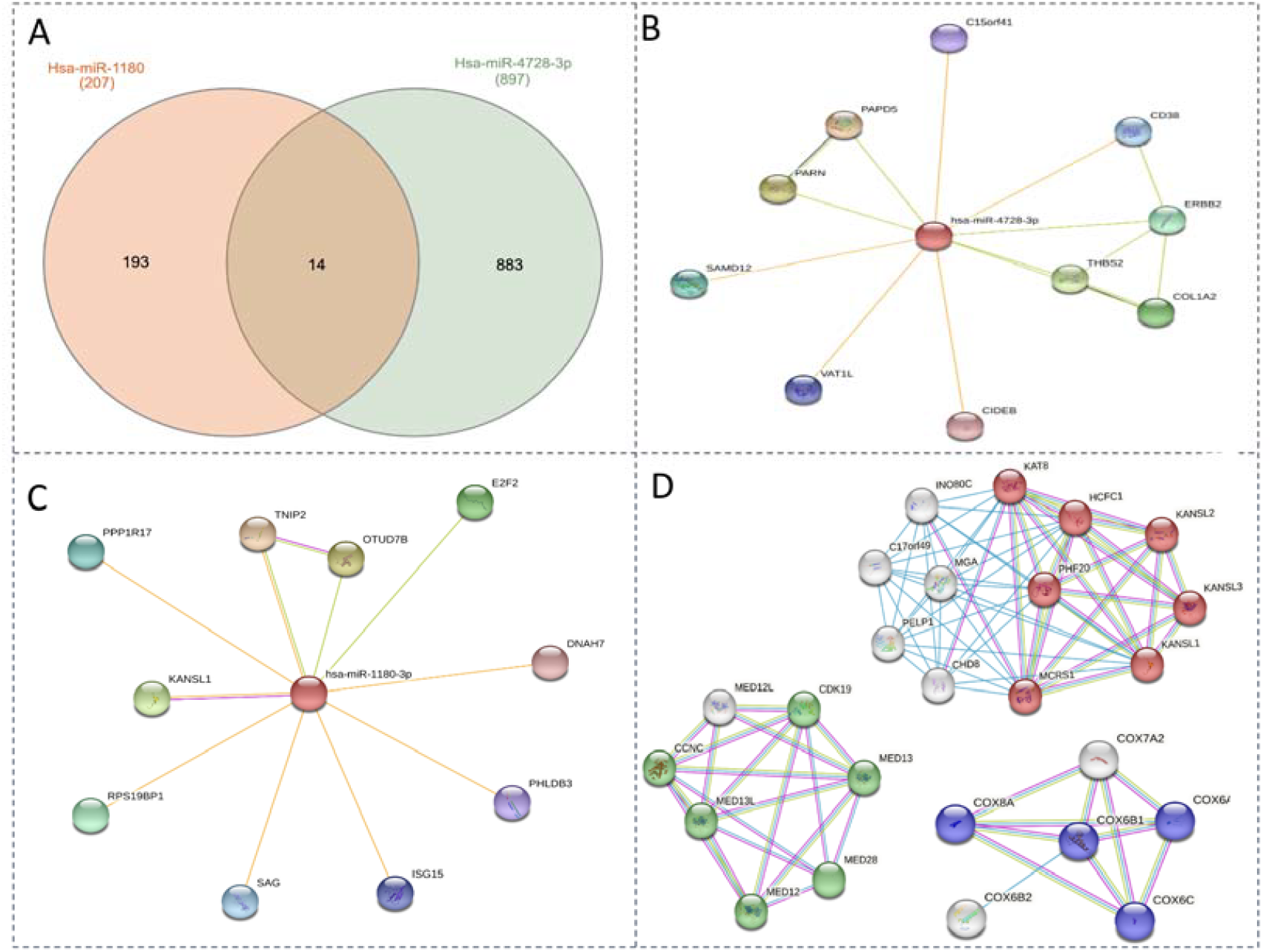
PPI and RNA-protein association analysis using STRING. (A) Venn diagram shows intersection of 14 miRNA targets between hsa-miR-1180 and hsa-miR-4728. (B, C) RNA–protein Association and Interaction Networks (RAIN) were identified. (D) Interaction of miRNA targets. STRING diagram of target PPI of hsa-miR-1180 and hsa-miR-4728 target proteins indicate potential role of epigenetic modifiers of HATs (red nodes), TP53 regulation of metabolic pathways (Blue nodes) and PPAR□ regulated gene expression in breast cancer (Green nodes). Lines indicate interaction evidence.

### Functional Impact on TNBC Progression

Our results demonstrate that miR-1180 and miR-4728, in particular, significantly enhance TNBC cell motility and invasiveness, as shown by wound healing and invasion assays. These miRNAs also share a subset of predicted targets involved in epigenetic regulation, TP53 signalling, and metabolic adaptation. The ability of these miRNAs to modulate both phenotypic traits and intracellular signalling underlines their functional role in sustaining the aggressive behaviour of TNBC. Of the five miRNAs, two were found to show exclusively high expression in TNBC clinical tissue. RT-qPCR analysis revealed a notable increase in the expression of hsa-miR-1180 and hsa-miR-4728 in TNBC tissues when compared to non-tumor breast tissues. Our study has confirmed that hsa-miR-1180 and hsa-miR-4728 act as tumor promoters, suggesting that their upregulation in tumor tissues may contribute to the progression and metastasis of TNBC. Decreased expression of these miRNAs attenuated tumor progression, as demonstrated by wound healing and invasion assays (Fig 7A, B)

### Comparison with Existing Literature

Previous studies have linked miR-6803, miR-1180, miR-4728, miR-1915, and miR-940 to various cancers, including colorectal, gastric, and cervical carcinomas. However, their roles in TNBC, particularly in the context of exosomal secretion and CSC biology, have not been characterized until now. Our study is the first to connect this miRNA panel to secretory pathways in TNBC and to validate their consistent expression across in vitro models and patient samples. Furthermore, our integrative approach reveals mechanistic convergence between exosomal miRNA signalling and known oncogenic pathways, a perspective largely absent in prior reports.

Using gene enrichment analysis and target prediction models, these dysregulated miRNAs were also found to be involved in a wide range of crucial biological processes and pathways including regulation of RNA pol II, Wnt signaling, Notch signaling, MAPK signaling pathway and the regulation of JNK signaling cascade. Previous studies have highlighted the correlation of Wnt signaling pathways with maintenance of stem cell niche and expression in cancer stem cells^35–41^. Notch receptor and ligand overexpression is linked to TNBC progression^38,42^. Notch receptors are also involved in modulating the behaviour of tumor-initiating cells and in the initiation of TNBC^38,42,43^. These pathways are pertinent to the therapeutic targeting of CSCs or other cells responsible for diverse prolapse as they play key roles in genesis and regulation of cell survival and their fate^44^. Given that many of these pathways are known to support tumor cell proliferation, cancer stem cell (CSC) survival, epithelial-to-mesenchymal transition (EMT), and invasion, it is likely that these miRNAs contribute to the progression and expansion of TNBC and TNBC stem cells.

Exosome-encapsulated miRNAs have a great potential as prognostic biomarkers. Several studies have demonstrated that exosomes are robust and may be stored for extended periods without compromising the integrity of encased miRNAs. These features greatly increase their potential applicability in a diagnostic or clinical setting. Secretory miRNAs are reflective of their parent cell status and thus may reveal a more specific tumor profile than conventional miRNA profile derived from whole blood or serum. From our identified set of five oncogenic miRNAs, miR 6803 has been previously identified as a diagnostic and prognostic marker in colorectal cancer, suggesting it may be a circulating marker for aggressive tumours. Yan et al found miR-6803-5p promotes tumor cell proliferation and invasion via the NF-κB pathway^54,55^., hyper-activation of which is often seen in TNBC. miR 1180 appears to be a potent oncomiR across multiple cancers and can activate Wnt pathway. Guo et al showed that plasma exosomal miR-1180-3p is a novel diagnostic marker for cutaneous melanoma. However, Tan et al. demonstrated that miR-1180 confers apoptotic resistance in hepatocellular carcinoma via NF-κB activation. This dual ability to activate Wnt and NF-κB signals suggests miR-1180 can promote survival, stemness, and therapy resistance, which might explain the aggressive behaviour of TNBC cells and stem-like cells expressing it. Studies have shown that miR-1180 from mesenchymal stem cells also promoted ovarian cancer cell glycolysis and proliferation via suppressing the Wnt inhibitor SFRP1. Similarly, miR-6803 has been shown to activate NF-κB signalling – in colorectal cancer, exosomal miR-6803-5p drove cancer cell proliferation and invasion via the PTPRO/NF-κB axis and could contribute to invasion or chemoresistance in TNBC.^56–61^ miR 1915 has been reported in gastric cancer^62,63^, colorectal carcinoma (CRC)^64–66^ and most recently in Breast cancer as a circulating biomarker for breast cancer (along with miR-455-3p)^67^. This demonstrates that miR-1915 can be detected in blood and has diagnostic value for breast malignancy. Interestingly, miR 4728 was found to have dual functions. It acts as a tumor suppressor in CRC^68^ while being tumorigenic in breast cancer^69–71^. miR 4728 has also been identified as a marker of HER2 status in BrCa patients. It is encoded within the HER2 gene, and although TNBC lacks HER2 amplification, recent studies in breast cancer highlight miR-4728’s oncogenic potential. Floros et al reported co-amplification of miR-4728 in HER2-positive breast cancers helps tumors evade anti-HER2 therapy – essentially, miR-4728 upregulation protected cancer cells from targeted drugs, underscoring its role in therapy resistance. While TNBC cells do not overexpress HER2, miR-4728 might still exert pro-tumor effects via HER2-independent pathways or by affecting other growth factor pathways; its presence in TNBC exosomes is therefore intriguing. miR 940 has been widely reported as an oncogenic marker for CRC^72^, gastric cancer^73^, cervical cancer^74^ and breast cancer^75,76^. Zhang et al demonstrated that miR-940 promotes breast cancer progression by downregulating the tumor suppressor FOXO3, indicating it can drive proliferation and invasion by silencing a key cell-cycle/ apoptosis regulator. This may explain the aggressive phenotype observed in TNBC cases with high miR-940. Rashed *et al* found that exosomal miR-940 from cancer cells can maintain oncogenic SRC signalling in recipient cells. It is involved in four critical pathways: the Wnt/β-catenin pathway, the MAPK pathway, the PD-1 pathway, and the phosphatidylinositol 3-kinase (PI3K)-Akt pathway, all of which are significantly implicated in breast carcinogenesis^77,78^. While that study was not in TNBC, it reinforces our hypothesis that tumor-derived exosomal miR-940 can alter the tumor microenvironment or distant sites to favour cancer progression

### Diagnostic and Therapeutic Potential

The secretory nature of these miRNAs and their detectability in exosomes make them attractive candidates for liquid biopsy-based monitoring. Their stability in circulation and association with tumor aggressiveness suggest they could serve as prognostic markers for early relapse or metastatic risk. Moreover, miR-1180 and miR-4728 may serve dual roles as biomarkers and therapeutic targets, given their contribution to invasive phenotypes. Targeted inhibition of these miRNAs could potentially disrupt key survival pathways and improve treatment responses. While many of these selected miRNAs have been identified in various types of cancer, their secretory roles in TNBC and TNBC stem cells (TNBCSCs) were previously unrecognized. Our study reveals the presence of secretory miRNAs (miRNAs miR 6803, miR 1180, miR 4728, miR 1915 and miR 940) in TNBC and TNBC stem cells highlighting their diagnostic value and clinical utility for better management of TNBC. Furthermore, these secretory oncomiRs were found consistently overexpressed in TNBC and TNBCSCs. We subsequently correlated our findings with TNBC tumor tissue samples (n=15) and found a consistent high expression of all five oncomiRs across tumor biopsies. From this panel of five miRNAs, two oncomiRs -hsa-miR-1180 and hsa-miR-4728 were found to be significantly upregulated across all tumor tissue samples in TNBC patients. The presence of these two secretory miRNAs in TNBC and TNBCSC along with their overexpression in clinical tissue samples indicate their possible role in TNBC progression and metastasis and may serve as reliable prognostic as well as therapeutic marker(s).

### Study Limitations and Future Directions

While our study presents a robust foundation, it is limited by the modest number of clinical samples analysed. Larger, multicentric cohorts will be necessary to validate the prognostic utility of the identified miRNAs. Functional studies in vivo are also warranted to confirm the role of these miRNAs in tumor progression and metastasis. Future research should explore combinatorial strategies involving miRNA inhibition and standard chemotherapy or immunotherapy to assess synergistic effects. Additionally, the integration of miRNA profiling with multi-omic datasets could provide a deeper understanding of TNBC heterogeneity and refine patient stratification strategies.

During the systematic literature review, the initial screening was conducted solely on three databases (PubMed, Web of Science and Cochrane Library), which may have resulted in the omission of some relevant studies. Ideally, we could have searched for additional data sources such as pre-print server archives, relevant books, Google Scholar and other similar resources. Additionally, the limited availability of publicly listed TNBC clinical data sets in GEO and TCGA may have narrowed the scope of our estimates. We have analyzed potential circulatory prognostic markers for TNBC, validated our findings in TNBC tumor tissues and further validated them in vitro using TNBC cell lines. Since TNBC is a highly metastatic disease, it is very challenging to obtain fresh surgical tissue samples. We were able to include a small number of tumor tissue samples which may limit the clinical relevance and translational impact of our study. We recognize that analysis of larger datasets and clinical correlation using higher numbers of TNBC tissues could provide greater insights on prognostic significance of these TNBC derived exosomal miRNAs. Further, addition of population-based studies and data stratification could provide better understanding and help in effective therapy and/or prognostic monitoring of TNBC.

Nonetheless, the identification of novel secretory miRNAs in TNBC and TNBCSCs is promising with potential application in the diagnosis and treatment of hormone refractory, metastatic breast cancers. Our study identifies novel oncogenic exomiRs in TNBC and TNBCSCs and highlights their utility as therapeutic targets for TNBC. The mechanisms through which these exomiRs influence the development and progression of TNBC are not yet fully elucidated. Nonetheless, it is believed that they may modulate the expression of genes associated with cell signalling pathways, including the Wnt, Notch, and MAPK pathways, which are known to play a role in the progression and relapse of TNBC. Previous studies have demonstrated similar prognostic potential of several other miRNAs in breast cancer such as miR-9^79^, miR-21^9^, miR-29b^9,34^ and miR 331^80^. Our study is the first to assess prognostic behaviour of these novel TNBC and TNBCSC derived exosomal miRNAs. Elucidating the role of secretory miRNAs in the molecular mechanisms driving TNBC initiation and progression is crucial for advancing miRNA-based therapeutics, which hold promise as effective treatment options for TNBC patients.

In conclusion, our study establishes a novel exosomal miRNA signature associated with TNBC and its stem-like subpopulations. These findings offer new avenues for biomarker development and therapeutic innovation, paving the way for more personalized and effective management of this challenging breast cancer subtype.

## Conclusion

Collectively, our findings underscore the potential of miR-1180, miR-4728, and their secretory counterparts as clinically relevant biomarkers for aggressive breast cancers. Future studies involving larger patient cohorts and longitudinal follow-up will be crucial to establish their utility in routine diagnostics or as therapeutic targets.This work opens a promising avenue for non-invasive miRNA-based monitoring and risk stratification in TNBC.

## Abbreviations

TNBC: Triple Negative Breast Cancer
miRNA: micro-RNA
TEM: transmission electron microscope
SEM: Scanning electron microscope
FBS: fetal bovine serum
DMEM: Dulbecco’s Modified Eagle Medium
GAPDH: glyceraldehyde 3-phosphate 26 dehydrogenase
PVDF: polyvinyl difluoride
RTqPCR: Reverse transcription-quantitative PCR
RIPA: Radio-Immunoprecipitation Assay
SDS: sodium dodecyl sulfate
PPI: Protein-Protein interactions
GO: Gene ontology
GEO: Gene expression omnibus
DEG: Differentially expressed genes

## Acknowledgements

This study was supported in part by the Department of Science and Technology (DST), Ministry of Science and Technology, Government of India by awarding the INSPIRE fellowship to A.Choudhary for her Ph.D. study. Nanoparticle Tracking Analysis was performed at the Translational Health Science and Technology Institute (THSTI), in Faridabad, India. We thank Ms Divya Arya (THSTI) for her guidance and expertise in handling Malvern NanoSight NS300 and exosome sample analysis. Confocal microscopy was performed at Regional Centre for Biotechnology (RCB) in Faridabad, India. We thank Mr Suraj Tewari for his technical expertise in Super-resolution microscopy and confocal imaging; Max Institute of Cancer Care, Max super specialty hospital, Saket, New Delhi for the provision of TNBC tissue samples for clinical validation; Mr. Akshey Kaushal, application engineer at the Indian Institute of Technology (IIT) Delhi, for his technical help with HR-TEM and Cryo-TEM employed for exosome visualization; Dr Prasanna Venkatraman, Deputy Director at the Cancer Research Institute, ACTREC Mumbai, India for sharing with us MCF10A cell lines; Mr. Manoj Gupta and Dr Pradeep K Rai for assistance with FACS analysis.

## Author Contributions

**B. C. Das:** Conceptualization, Supervision, Reviewing and Editing; **Ananya Choudhary:** Data curation, Methodology, Investigation, Writing, Original draft preparation, Reviewing and Editing; Funding acquisition; **Satish S. Poojary**: Visualization, Methodology, Supervision; **Priyanka Jain**: Software, Validation, Formal Analysis**; Harit Chaturvedi :** Resources

## Availability of supporting data

All data relevant to the study are included in the article or uploaded as supplementary information.

## Declarations

### Ethics approval and consent to participate

Informed consent was obtained from all participants prior to their involvement in the study, and all procedures were conducted in accordance with the ethical guidelines of Indian Council of Medical Research (ICMR). Participants were fully informed about the study objectives, potential risks and benefits, and their right to withdraw at any time. Written consent forms were obtained from all participants, ensuring their privacy and confidentiality of data collected. All personal data and biopsy samples were handled with strict confidentiality. Identifiable information has been removed or anonymized in all published findings. All clinical samples were collected with signed informed consent and ethics approval also obtained from Max Institute of Cancer Care, Max Hospital Saket, New Delhi

### Consent for publication

The content of this paper has not been submitted to any other scientific publications. All the authors have declared that no financial conflict of interest exists. All authors have approved the submission of this work for publication in Breast Cancer Research.

