## Supplementary file 1 for "Identification of novel exosomal miRNAs and their role in diagnosis and prognosis of Triple Negative Breast Cancer"

Prof. Bhudev C. Das,

Amity Institute of Molecular Medicine & Stem Cell Research (AIMMSCR),

Amity University, Sector-125, Noida, 201313,

Uttar Pradesh, India.

**Supplementary Figures : Workflows and Experimental Strategies**

**Figure 1**

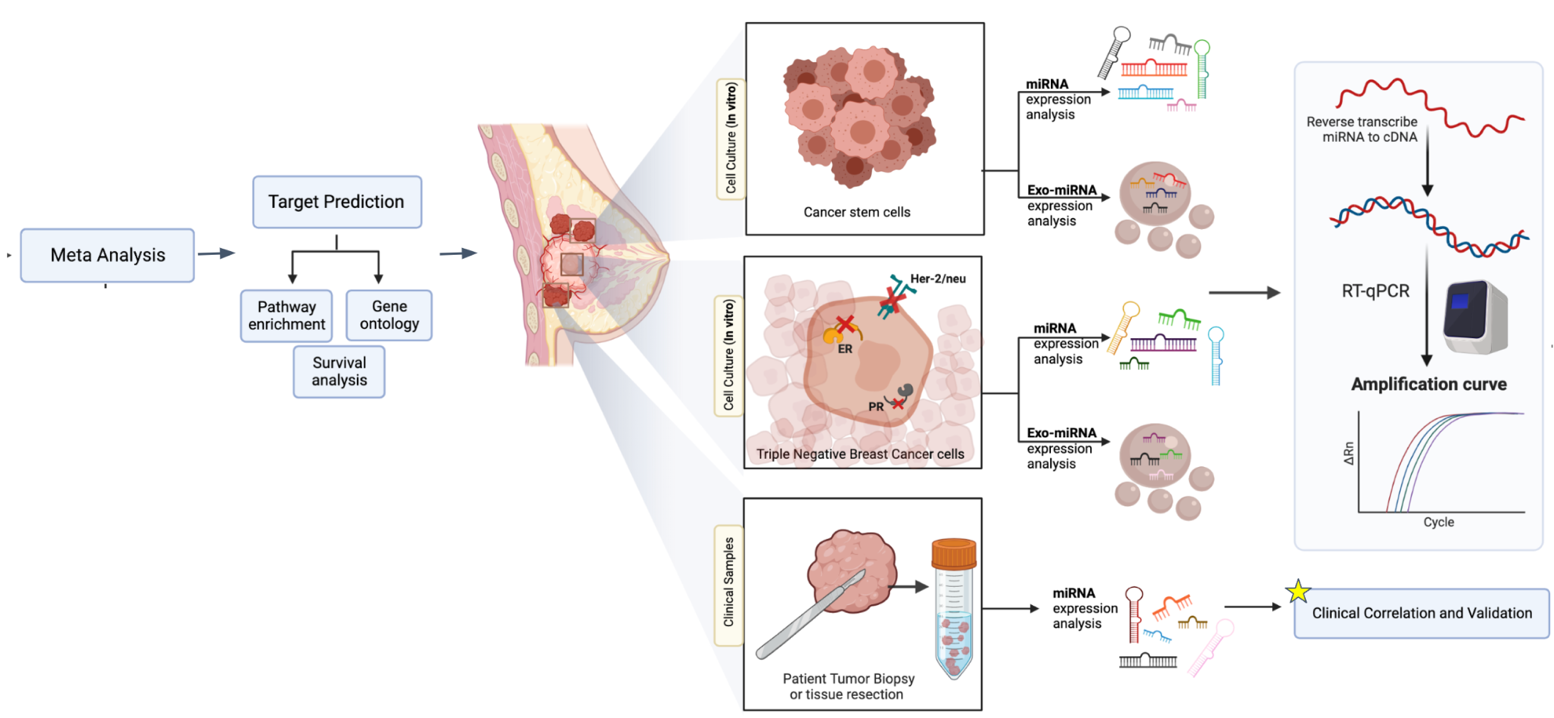

S1(Fig 1):  Study workflow and etiological model for analysis of exomiRs released by TNBC and TNBC stem cells; figure created with Biorender.com

**Figure 2**

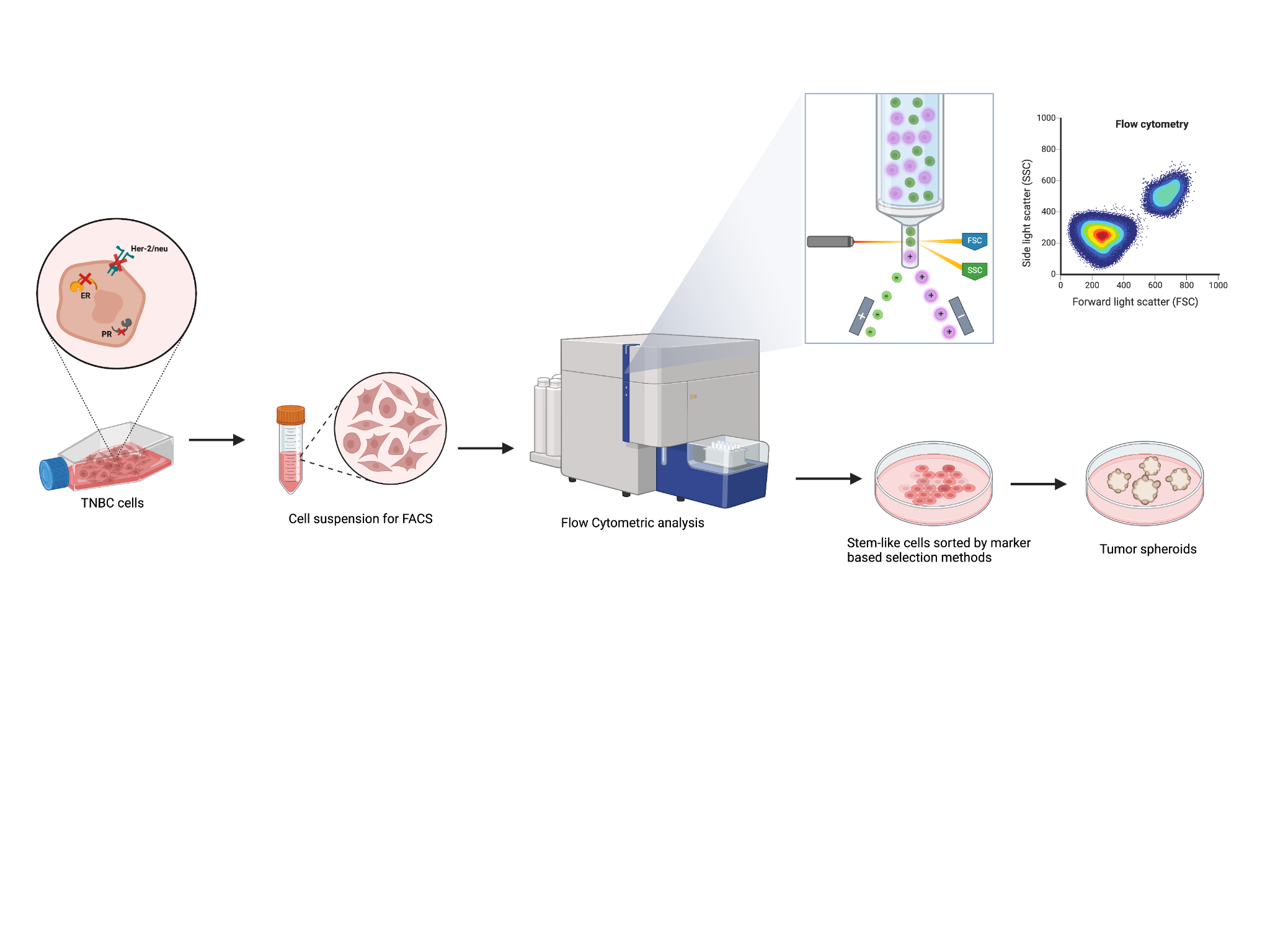

S1 (Fig 2) Representative model (workflow) depicting the isolation and characterization of TNBC derived stem-like cells. Cancer cell suspension is analysed by fluorescence activated cell sorting machines (FACS) by means of marker-based selection methods and dye-exclusion assays. Figure created with Biorender.com

**Western Blots**

**WB Full size images**

Full size blots for Fig 4 (d): Immunoblot showing expression of CD81 exosomal marker in TNBC cell line MDA-MB 231 derived exosome samples. Calnexin is used as a negative control while β-Actin is used as a positive control

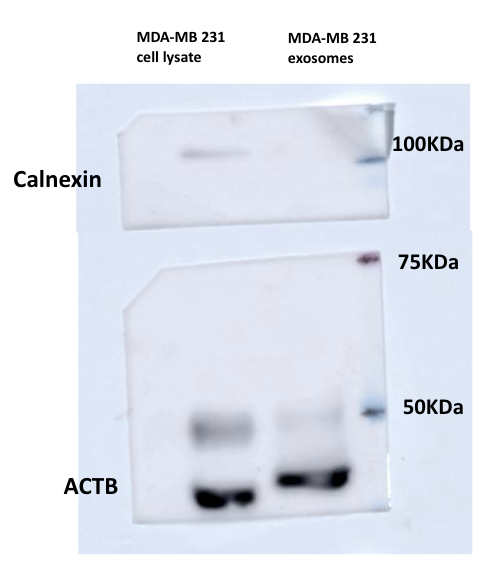

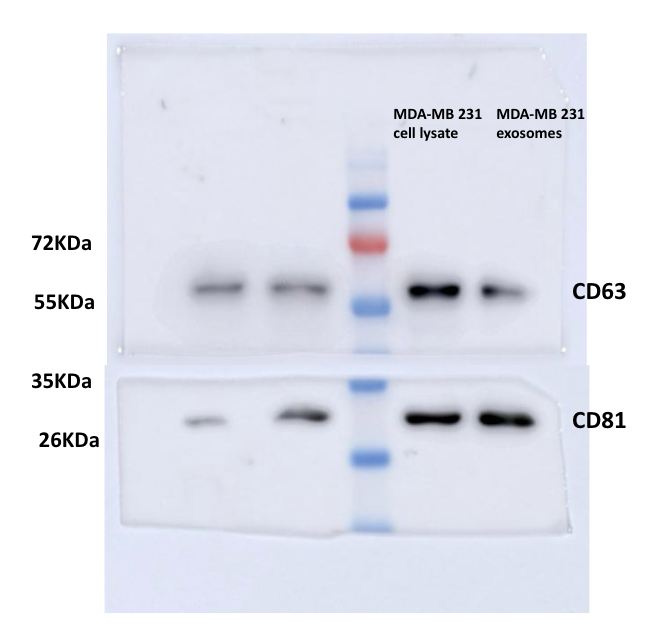

Full size blots for Fig 6 (H): Immunoblot showing expression levels of SOX2 in TNBC tumorspheres and cancer cells. GAPDH was used as positive control

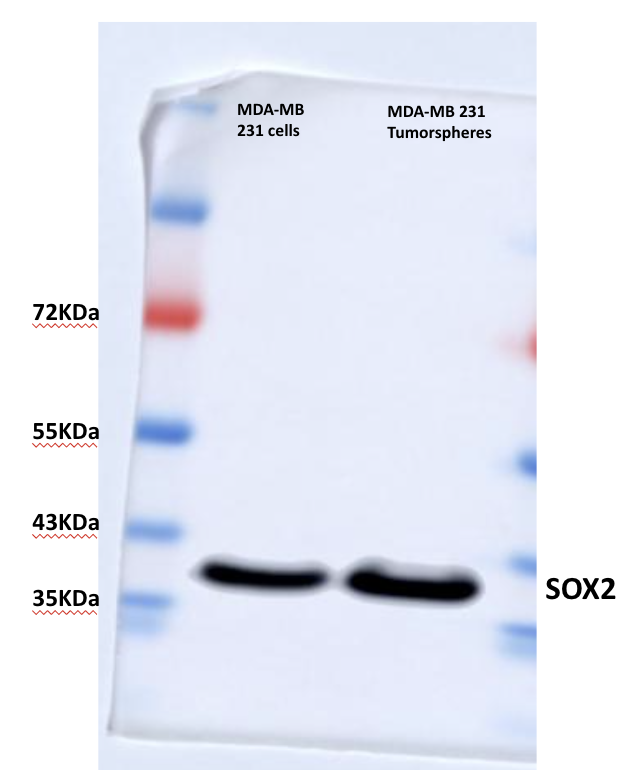

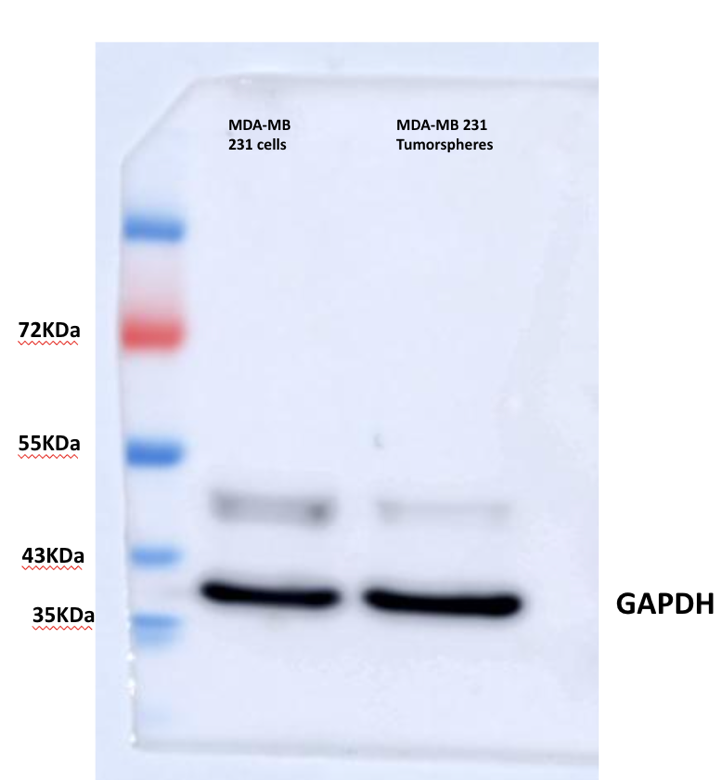

**Supplementary Table 1 (S1) : List of RT-qPCR primers used in the study**

| SNo. | Primers | Sequence |
| --- | --- | --- |
| 1 | **Sox2** | F: CGAGATAAACATGGCAATCAAATG  R: CACCAGAACAAATTCCGTTTGCAA |
| 2 | **ABCG2** | F: GCAGATGCCTTCTTCGTTATG  R: TCTTCGCCAGTACATGTTGC |
| 3 | **ALDH1** | F: GTCCTACTCACCGATTTGAA  R: CTTGTATAATAGTCGCCCCC |
| 4 | **miR 1915-3p**  (stem loop) | GTTGGCTCTGGTGCAGGGTCCGAGGTATTCGCACCAGAGCCAACCCCGCC |
| 5 | **miR 1180-3p** (stem loop) | GTTGGCTCTGGTGCAGGGTCCGAGGTATTCGCACCAGAGCCAACTATTCC |
| 6 | **miR 940**  (stem loop) | GTTGGCTCTGGTGCAGGGTCCGAGGTATTCGCACCAGAGCCAACACCACA |
| 7 | **miR 6803**  (stem loop) | GTTGGCTCTGGTGCAGGGTCCGAGGTATTCGCACCAGAGCCAACACGCCC |
| 8 | **miR 4728**  (stem loop) | GTTGGCTCTGGTGCAGGGTCCGAGGTATTCGCACCAGAGCCAACCTGGGG |
| 9 | **miR 1180-3p** (Forward) | TTTTTGGACCCACCCGGC |
| 10 | **miR 1915-3p**  (Forward) | TTTTTCCCCAGGGCGACG |
| 11 | **miR 940**  (Forward) | TTGTTAAGGCAGGGCCCC |
| 12 | **miR 6803**  (Forward) | TTTTTCTGGGGGTGGGGG |
| 13 | **miR 4728**  (Forward) | GTTTCATGCTGACCTCCCT |
| 14 | **Universal Reverse** | GTGCAGGGTCCGAGGT |
